# Interrogating the Mechanisms of Cas9-mediated Allele Conversion

**DOI:** 10.64898/2026.02.17.705987

**Authors:** Joss B. Murray, Emma Collins, Lisa Lonetti, Lucia Nicosia, Tadhg Crowley, Ciaran M. Lee, Patrick T. Harrison

## Abstract

Allele conversion describes a process where a heterozygous variant is made homozygous. Recently, it has been shown that allele conversion can be triggered by DNA damage at the heterozygous site. This process has the potential to repair pathogenic heterozygous mutations; however, the efficiency is low. Here, we endeavoured to understand the mechanism underlying allele conversion, ultimately to raise allele conversion efficiency to functionally relevant levels. To test this, we developed a Compound Heterozygous Allele Conversion Reporter (CHACR) cell line. This line comprises knocked-in fluorescent protein encoding genes, with heterozygous inactivating mutations resulting in different fluorescence profiles from each allele. These mutations create protospacer adjacent motifs (PAM) for Cas9 recognition, where allele-specific gRNAs (AS-gRNAs) target the heterozygous mutations. We showed that applying these AS-gRNAs with either Cas9 nuclease or Cas9(D10A) nickase can recover mCherry fluorescence. Sorting and sequencing these fluorescent cells revealed wild-type sequences, suggesting allele conversion repaired the mutation using the homologous allele as a template. Allele conversion can also be triggered using an adenine base editor with an AS-gRNA, and this allele conversion mechanism can be manipulated by inhibiting DNA-PKcs or overexpressing RAD51. This work introduces a model for measuring allele conversion, and modifiers of this mechanism.

## Introduction

The underlying cause of genetic disease stems from mutations which change the nucleotide sequence. In the case of a recessive genetic disease such as cystic fibrosis, both copies of the gene must be mutated to produce the pathogenic phenotype. Modern therapeutic strategies for genetic diseases fall under two categories: gene therapies and gene editing. A recent example of this is the approval of two different treatments for Sickle Cell Disease (SCD), one is a classical gene therapy (Lyfgenia) which involves the delivery of an exogenous cDNA to produce the missing protein to ameliorate the disease symptoms. The other is gene editing which modifies the endogenous genomic material in the cell in a targeted manner. In the case of SCD, the clinically approved strategy has been to use editing to disrupt BCL11A which encodes a transcriptional regulator thus enabling the expression of fetal hemaglobin to compensate for this missing gene (Leonard and Tisdale, 2024). Other strategies include base editing (Newby et al., 2021), HDR and prime editing (Everette et al., 2023). In the case of SCD, one specific mutation accounts for nearly all manifestations of SCD (Tsukahara et al., 2024), meaning that one editing strategy has wide reaching potential. However, for many other genetic diseases, the wide range of pathogenic variants means multiple editing strategies must be devised.

With the increasing innovation on these gene editing technologies, novel strategies are improving the efficiency at the cost of complexity. Additionally, these new strategies often require more and larger components making clinically relevant delivery challenging. Therefore, there is a niche for identifying methods that can correct pathogenic mutations that are less cumbersome. HDR offers some flexibility for gene editing, although this strategy requires a donor with the specified corrective sequence and also risks indels from the required Cas9 nuclease. However, in the case of dominant genetic diseases caused by a single faulty allele, or recessive diseases caused by compound heterozygous mutations, the requisite corrective sequence naturally exists on the homologous allele.

Recent evidence has suggested the potential of CRISPR-based systems to create allele-specific DNA damage in a heterozygous genotype, leading to repair of the genetic lesion using information derived from the homologous allele thereby negating the need for an exogenous donor sequence (Wilde et al., 2021, Roy et al., 2022, Tomita et al., 2023, Yang et al., 2023). This concept depends on the sequence targeted for conversion being heterozygous, which is often the case for recessive diseases with multiple causative variants. This also requires the mutation to create a site that will be targeted by a nuclease on one allele but not on the other. For example, Cas9 requires a protospacer adjacent motif (PAM) with an NGG sequence for optimal binding. Therefore, a heterozygous variant destroying or creating a PAM sequence will result with allele-specific Cas9 binding site. With this recent evidence of allele-specific, Cas9 mediated DNA damage triggering allele conversion, we aimed to investigate the mechanism surrounding this process.

We devised a fluorescence reporter cell model, where genes encoding fluorescent proteins in tandem include heterozygous, inactivating mutations. We show that guide RNAs (gRNAs) targeting these mutations have the capacity trigger allele conversion and copy the information from the other allele, leading to mutation correction and observable fluorescence output. Using this model, we show that these allele-specific gRNAs can trigger allele conversion and induce fluorescence using either a Cas9 nuclease or a Cas9(D10A) nickase. Furthermore, we investigate the mechanism behind allele conversion by modifying factors involved in DNA repair, illustrating the potential of existing molecular processes to trigger corrective interchromosomal recombination.

### Design of a Compound Heterozygous Allele Conversion Reporter (CHACR) cell line

To create a compound heterozygous fluorescence reporter cell line to study Cas9-mediated allele conversion, we first designed two near-identical plasmids (B and G) encoding either wild-type or mutant versions of three different fluorescent proteins (EGFP, mCherry and mTagBFP2). The constructs, which differ by just 9nt across a ~6 kb region, were flanked by homology arms to enable site-specific integration into AAVS1 safe-harbor locus of chromosome 19 in HEK293T cells (Figure 1A).

**Figure 1.**
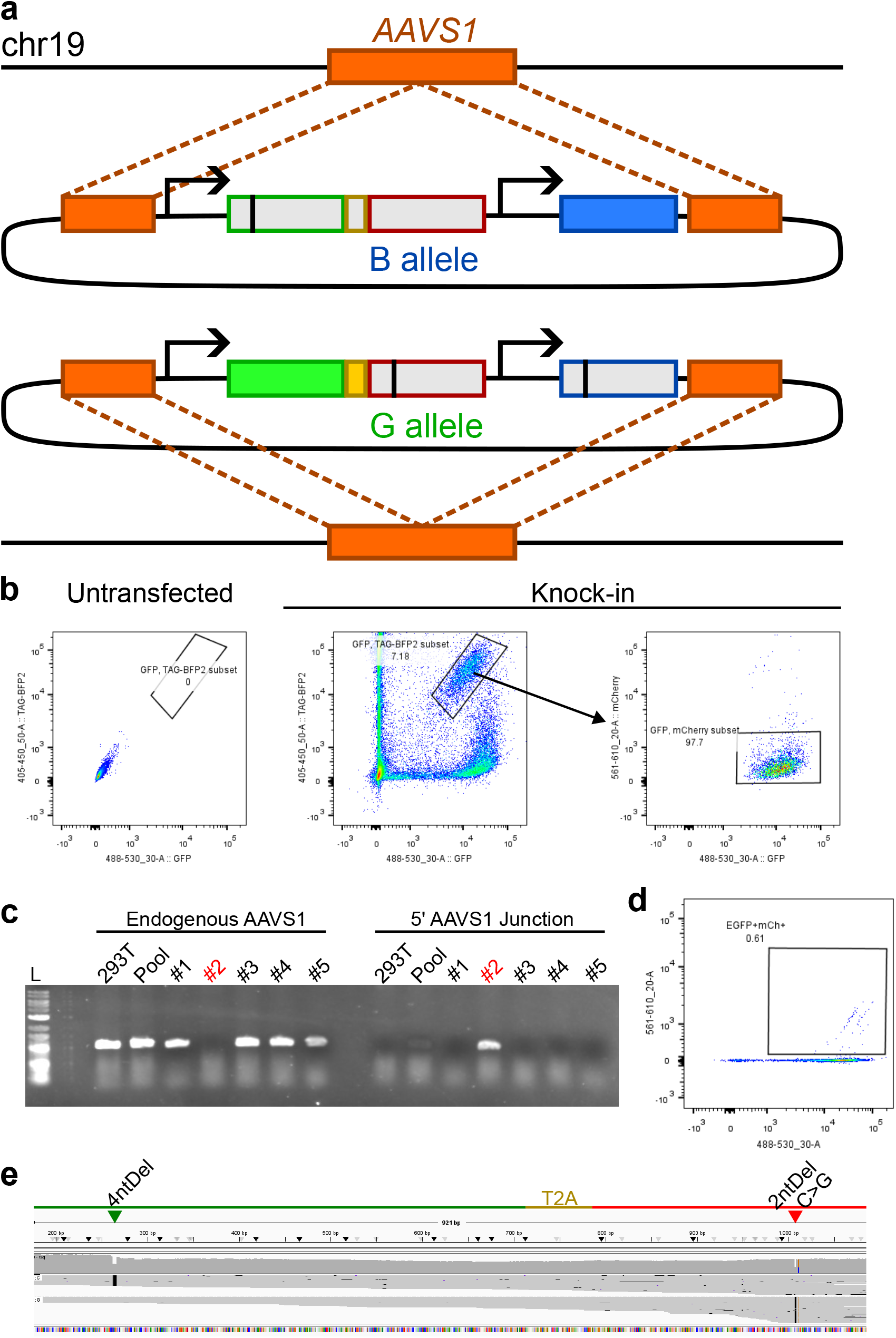
(a) Schematic of the B and G allele expression plasmids, and integration into the *AAVS1* locus. Each allele is comprised of a CAG promoter driving the expression of EGFP linked to mCherry with a T2A self-cleaving peptide. Downstream, a CMV promoter drives the expression of mTagBFP2. On the B allele, a 4nt deletion disrupts the reading frame of EGFP and mCherry, leading to only mTagBFP2 expression from this allele. On the G allele, a PTC mutation in mCherry and a 2nt insertion in mTagBFP2 leads to only EGFP expression from this allele. (b) Flow cytometry gating strategy for sorting cells expressing both EGFP and mTagBFP2. Untransfected HEK293T cells express neither EGFP or mTagBFP2, while potential knock-in clones that are positive (GFP+ TAG-BFP2 subset) are gated, and are also mCherry negative. (c) Diagnostic PCR for the endogenous *AAVS1* locus as well as the 5’ *AAVS1* knock-in junction. (d) Flow cytometry confirmation of EGFP and mCherry fluorescence from CHACR clone #2. (e) IGV visualisation of alignment between a reference sequence with wild-type EGFP and mCherry against a PCR amplicon of the EGFP and mCherry cassette amplified from CHACR clone #2 confirming the heterozygosity and position of each fluorophore inactivating mutation.

The B allele plasmid comprises a CAG promoter driving EGFP with a 4nt deletion (EGFP^Δ4^) linked to wild-type mCherry via a T2A sequence. The 4nt deletion creates a frameshift (fs) which inactivates EGFP, which also shifts mCherry out of frame leading to no expression. This plasmid also contains a CMV promoter which expresses a functional mTagBFP2 reporter. When integrated into cells, this allele can only express a functional blue fluorescent protein, hence the designation as the B (blue) allele. This design allowed us to create an allele-specific gRNA (AS-gRNA) that targets the EGFP^Δ4^ in the integrated B allele. If transfection of this AS-gRNA with Cas9 nuclease or nickase causes editing of the mutation by allele conversion, then this will result in expression of a functional EGFP protein from the B allele. A more important feature of our design is that successful allele conversion will result in wild-type mCherry expression as the T2A sequence is also now in-frame, and this will allow the isolation of mCherry positive cells for sequence analysis.

The G allele plasmid comprises a CAG promoter and a wild-type EGFP reporter linked to a mutant mCherry^fs-2;Y77X^ via a T2A sequence. The mCherry^fs-2;Y77X^ has a 2nt deletion (CC) which creates a frameshift (which inactivates mCherry) and a C>G SNP (Y77X) which creates an NGG PAM site. This plasmid also contains a CMV promoter and a variant of the mTagBFP2^fs+2^ reporter which is non-functional due to a 2nt insertion (GG), also creating an NGG PAM site. When integrated into cells, this allele can only express a functional green fluorescent protein and is thus called the G (green) allele. This design allowed us to create an AS-gRNA that targets the mCherry^fs-2;Y77X^ in the integrated G allele. If transfection of this AS-gRNA with Cas9 nuclease or nickase results in editing of the mutation by allele conversion, then this will result directly in expression of a functional wild-type mCherry protein and allow the isolation of mCherry positive cells for sequence analysis.

To make the cell line, the B and G allele plasmids were co-transfected into HEK293T cells with plasmids encoding Cas9 nuclease and an *AAVS1*-targeting gRNA. To identify cells which had both allele sequences stably integrated into the *AAVS1* loci of the homologous chromosome 19 sites, fluorescence-activated cell sorting (FACS) was performed to select single cells positive for both green and blue fluorescence (Figure 1B). A potential clone (#2) was initially characterised by PCR as positive for the 5’ *AAVS1* junction bordering the CAG promoter and negative for the endogenous *AAVS1* locus (Figure 1C). Further analysis of this clone by ONT sequencing confirmed these cells were compound heterozygous for the knocked-in B and G alleles. As shown in Figure 1C, the 4nt deletion in EGFP^Δ4^ on the B allele was in phase with a wild-type mCherry sequence, and a wild-type EGFP on the G allele was in phase with mCherry^fs-2;Y77X^ harbouring a 2nt deletion and C>G SNP followed by a Y77X PTC mutation. This clonal line of cells showed a 1:1 heterozygosity of B allele to G allele (Figure S1) and is subsequently referred to as the Compound Heterozygous Allele Conversion Reporter (CHACR) clone. Another heterozygous clone was identified with a 3:1 ratio of B:G alleles, suggesting that the parent line may be tetraploid (Supplementary Table 1), which is discussed later.

### Use of mCherry^fs-2;Y77X^ AS-gRNA with either Cas9 Nuclease or Cas9(D10A) Nickase to Target mCherry^fs-2;Y77X^ in the G allele Results in mCherry Fluorescence via Allele Conversion

The design of the B and G alleles allowed for AS-gRNAs to target the fluorescence-inactivating mutations (Figure 2A). CHACR cells were transfected with these AS-gRNAs and either Cas9 nuclease or Cas9(D10A) nickase, followed by flow cytometry. We used flow cytometry to identify the proportion of mCherry postive cells, 72h post-transfection (Figure 2B). Gating conditions were set such that ≤0.6% of non-transfected (Ctrl) and mock-transfected (Tfxn) reporter cells were scored as mCherry-positive (Figure 2C). To determine the genetic basis for the rescue of mCherry expression, we harvested genomic DNA from these cell populations and performed Illumina short-read NGS for two separate amplicons, one spanning the mCherry cassette (Figure 2D), the other for the EGFP cassette (Figure 2E). We classified these amplicons into one of four read categories as described in Table 1. As expected, analysis of both the EGFP and mCherry amplicons from the Tfxn only control cell populations showed a 50:50 ratio of B:G allele reads with no additional sequence changes observed.

**Figure 2.**
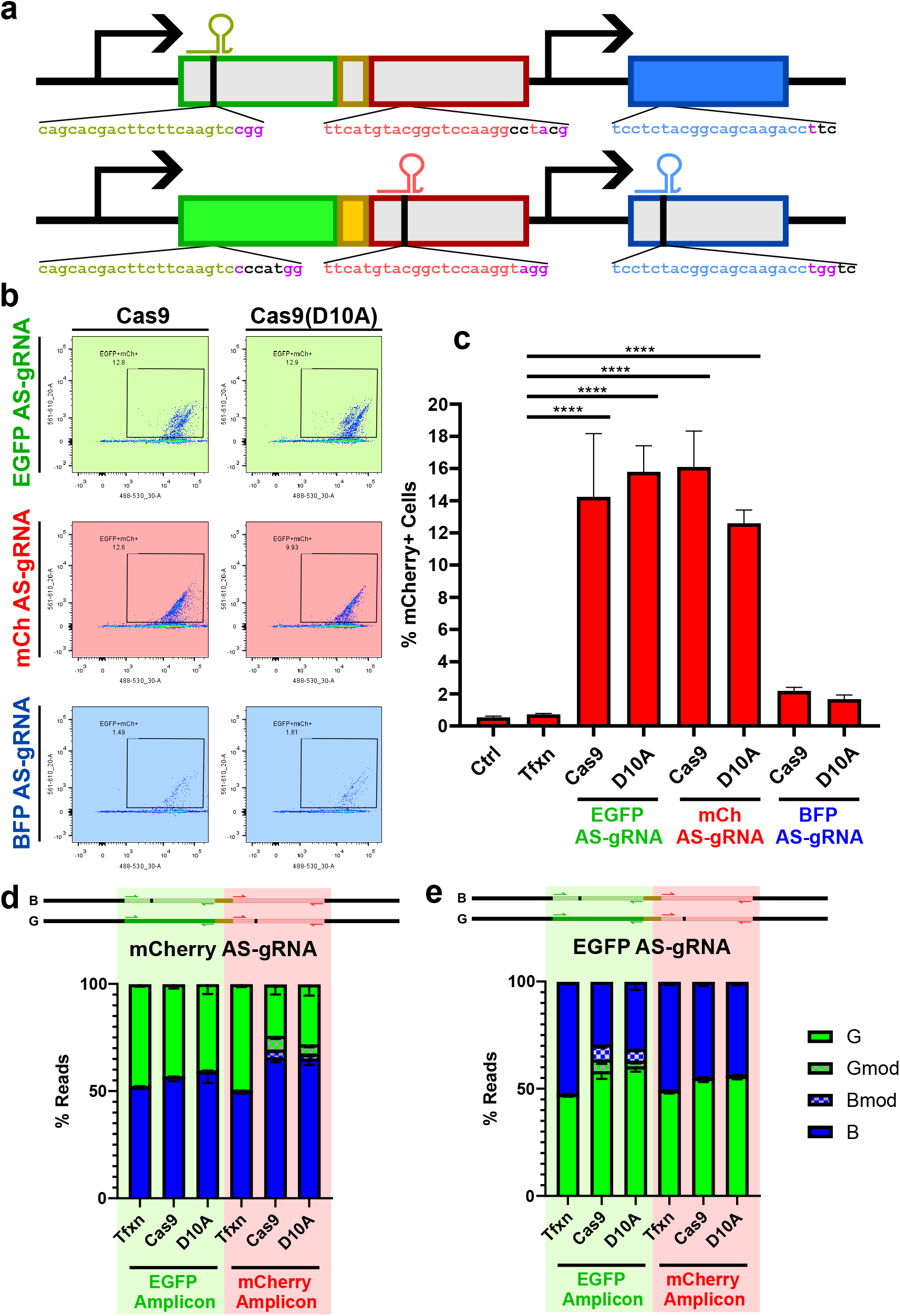
(a) Positions of AS-gRNAs targeting the fluorophore inactivating mutations across each allele. (b) Dot plots from CHACR cells transfected with Cas9 nuclease or Cas9(D10A) and each AS-gRNA. (c) Quantification of mCherry fluorescent cells highlighted within gates from (b). (d,e) Graphical representation of PCR amplicons produced for the EGFP, or mCherry, sequences after transfection with Cas9 or Cas9(D10A) and either the (d) mCherry or (e) EGFP AS-gRNAs. Resulting amplicons were sequenced by Illumina NGS 72h after transfection and analysed using CRISPResso2 for allele sequences matching either the B allele or G allele for the sequences amplified. Bars represent mean values from n=3 with error bars ± SD. *p* values calculated by one way ANOVA with Dunnett’s multiple comparisons posthoc test. * = *p*<0.05, ** = *p*<0.01, *** = *p*<0.001, **** = *p*<0.0001.

**Table 1:** Descriptive annotations of read nomenclature.

| Read Nomenclature | mCherry Amplicon | EGFP Amplicon |
| --- | --- | --- |
| **B allele** | Wild-type mCherry sequence | EGFP<sup>Δ4</sup> sequence |
| **B allele modified** | Wild-type mCherry sequence *plus* additional sequence changes | EGFP<sup>Δ4</sup> sequence *plus* additional sequence changes |
| **G allele** | mCherry<sup>fs-2;Y77X</sup> sequence | Wild-type EGFP sequence |
| **G allele modified** | mCherry<sup>fs-2;Y77X</sup> sequence *plus* additional sequence changes | Wild-type EGFP sequence *plus* additional sequence changes |

When CHACR cells were transfected with the mCherry^fs-2;Y77X^ AS-gRNA and either Cas9 nuclease or Cas9(D10A) nickase, we observed 16.1% or 12.6% mCherry postive cells, respectively (Figure 2B,C), suggesting repair of the mCherry^fs-2;Y77X^ sequence on the G allele. This was confirmed upon analysing the sequence of the mCherry amplicons which showed a ~15% increase in B allele mCherry reads (Figure 2D). This observation is consistent with a repair mechanism mediated by short allele conversion tracts where part of the wild-type mCherry sequence from the B allele acts as a template for repair of the mCherry^fs-2;Y77X^ sequence on the G allele to the wild-type and now functional mCherry sequence. We also noted an ~8% increase in B allele reads in the EGFP amplicons in these cells. The detection of such reads indicates an increase in the number of cells with EGFP^Δ4^ sequences, an observation consistent with a repair mechanism mediated by allele conversion, but with a longer editing tract that initiates at the mCherry^fs-2;Y77X^ site and extends into the EGFP^Δ4^ sequence on B allele such that it acts as a template for install the EGFP^Δ4^ sequence in the wild-type EGFP sequence. Of note, we also detected a number of reads referred to as “B allele modified” or “G allele modified”, in the mCherry amplicons, the relevance of which are discussed later.

### Use of the EGFP^Δ4^AS-gRNA with either Cas9 nuclease or Cas9(D10A) nickase to target EGFP^Δ4^ in the B allele also results in mCherry Fluorescence via allele conversion

Our TLR design also allowed us to test if mCherry expression could be restored by targeting the EGFP^Δ4^ in the B allele as precise repair of the 4nt deletion in EGFP^Δ4^ would bring the wild-type mCherry sequence back into the correct reading frame for translation. When cells were transfected with either Cas9 nuclease or Cas9(D10A) nickase plus the EGFP^Δ4^ AS-gRNA, a similarly large proportion of mCherry positive cells were observed, 14.2% and 15.8%, respectively (Figure 2B,C) suggesting precise repair of the B allele EGFP^Δ4^ had occurred, and that this restored the reading frame which enable expression of wild-type mCherry from the B allele.

This was confirmed upon analysing the sequence of the mCherry amplicons which showed a ~12% increase in G allele reads (Figure 2E). As shown in the diagram below the graph, this observation is consistent with a repair mechanism mediated by short allele conversion tracts where part of the wild-type EGFP sequence from the G allele acts as a template for repair of the EGFP^Δ4^ sequence on the B allele. This restores the reading frame and wild-type mCherry is now expressed from the B allele. A substantial proportion of the EGFP amplicons reads were identified as “B allele modified” or “G allele modified” (Figure S2A-D), the relevance of which is discussed later.

When we analysed mCherry amplicons in these cells, we observed smaller ~6% increase in G allele reads. The detection of such reads indicates an increase in the number of cells with mCherry^fs-2;Y77X^ sequences, an observation consistent with repair by allele conversion with a longer editing tract that initiates at the EGFP^Δ4^ AS-gRNA target site and which extends into the mCherry sequence on B allele and serves as a template to install the mutant mCherry sequence.

Application of the mTagBFP2^fs+2^ AS-gRNA did not significantly increase the number of mCherry-positive cells above baseline (Figure 2B,C). The resulting mCherry positive population was too small to sort, therefore subsequent NGS was not performed on CHACR cells after use of the mTagBFP2^fs+2^ AS-gRNA. This low level of mCherry fluorescence may be evidence of a directional or distance component required for this allele conversion mechanism. Together, these findings suggest that allele-specific DNA damage at the site of a mutation can shift the ratio of sequences towards that found on the other allele, and that this is associated with functional recovery.

### Long-read Sequence Analysis of mCherry Positive Cells Confirms Allele Conversion

Whilst our short-range PCR and Illumina sequencing data (Figure 2) strongly suggest that repair is occurring by an allele conversion mechanism, a notable limitation of this approach reads is that we can only sample one portion of the B and G alleles at any one time. Whilst this allows us to infer the generation of “wild-type” alleles, that is the existence of wild-type EGFP and wild-type mCherry on the same allele, this approach does not allow us to determine this directly. To address this, we sorted cell populations expressing both EGFP and mCherry (EGFP+/mCh+) shown in Figure 2B that were obtained from transfections with Cas9 or Cas9(D10A) nickase and either the EGFP^Δ4^ or mCherry^fs-2;Y77X^ AS-gRNAs, and harvested genomic DNA. We then generated long PCR amplicons spanning the beginning of EGFP to the end of mCherry and analysed them by Oxford Nanopore Technology (ONT) sequencing, phasing, and allele proportion as shown in Figure 3A.

**Figure 3.**
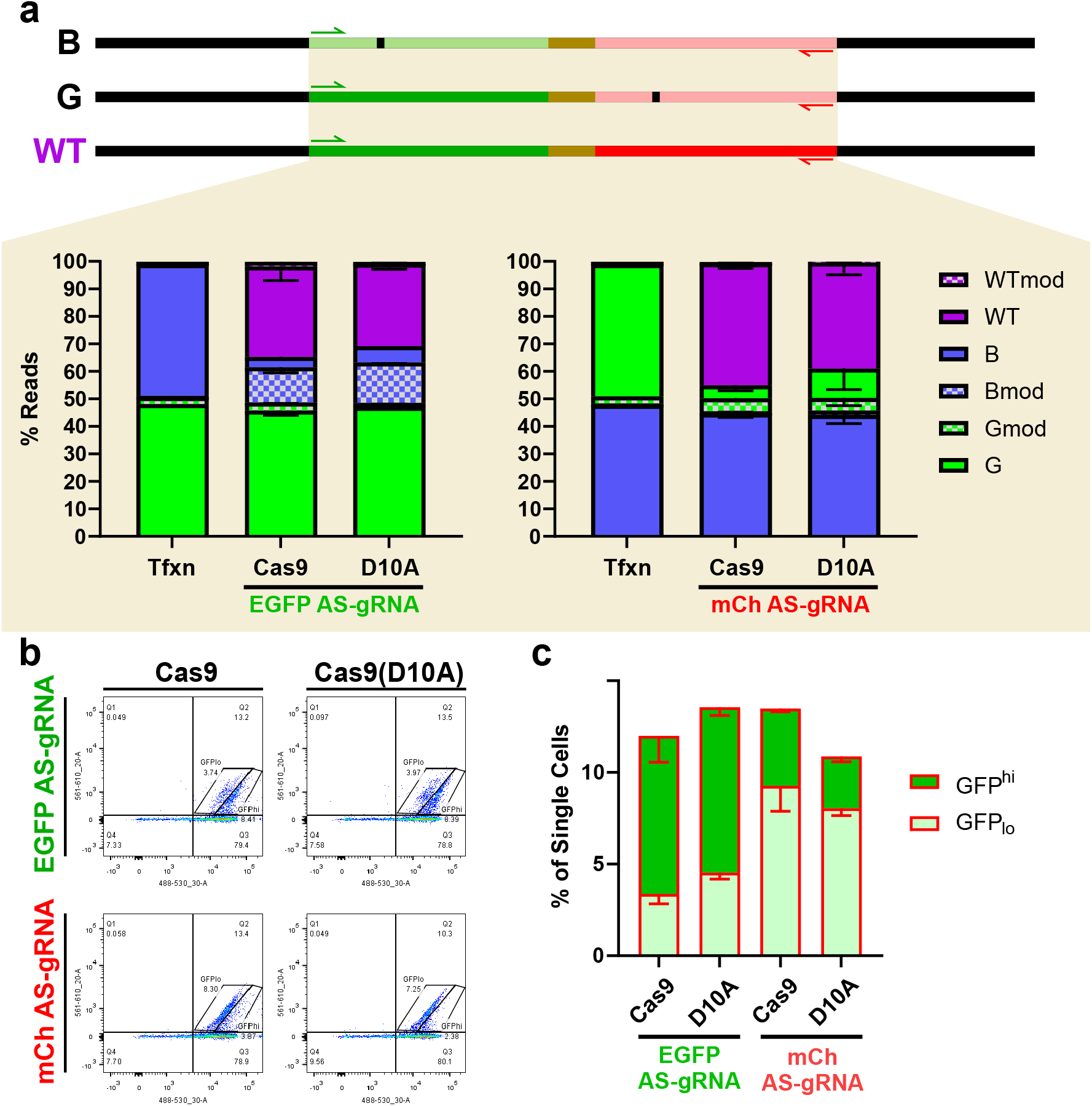
(a) Graphical representation of PCR amplicons for the EGFP and mCherry sequences on the B allele, G allele, and a potential wild-type (WT) allele that may arise after allele conversion. This PCR was conducted on CHACR cells sorted for mCherry fluorescence 72h after transfection, and sequenced by ONT. Sequencing data was analysed by CRISPResso2 for allele sequences matching either the B allele, G allele or a potential WT allele. (b) Dot plots from CHACR cells transfected with Cas9 nuclease or Cas9(D10A) and either EGFP or mCherry AS-gRNAs. Gates represent the EGFP high expressing (GFP^hi^) or EGFP low expressing (GFP_lo_) cells. (c) Quantification of GFP^hi^ and GFP_lo_ cells as subsets of the total mCherry fluorescent population after allele conversion. Bars represent mean values from n=3 with error bars ± SD.

In the EGFP+/mCh+ population of cells obtained following transfection with the EGFP^Δ4^ AS-gRNA plus either a Cas9 nuclease or Cas9(D10A) nickase, ONT sequence analysis showed almost no change in G reads as expected. However, more than half of the B alleles were converted to wild-type alleles (Figure 3A). The majority of remaining B reads now scored as B^mod^; close analysis of these modified reads showed that the most frequent modified B allele read is a +1nt insertion (Figure S3A-D). This insertion would restore the reading frame from −4nt into −3nt, thereby leading to mCherry expression (and thus selection of this population). We cannot rule out that these are imperfect allele conversion events, but a much more likely explanation is that these are simply the +1nt insertions mediated by an NHEJ mechanism which is commonly reported for both Cas9 nucleases and Cas9 nickases. Of note, a small proportion of B alleles remained; at first glance this is unexpected as the B allele does not permit the expression of a functional mCherry, so these cells should not have been selected. However, as noted earlier, the CHACR cells are tetraploid with two B alleles and two G alleles, thus these reads are most likely to be derived from cells where one B allele is converted to WT, but the other remains unchanged (see Supplementary Table 1).

In the EGFP+/mCh+ population of cells obtained following transfection with the mCherry^fs-2;Y77X^ AS-gRNA plus either a Cas9 nuclease or Cas9(D10A) nickase, almost no change in B reads was observed. However, most of the G alleles were converted to wild-type alleles (Figure 3A). Of note, there were far fewer modified reads than when using the EGFP^Δ4^ AS-gRNA. This is not to say that there is a substantial difference in indel formation when using the mCherry^fs-2;Y77X^ AS-gRNA; rather that a simple indel is unlikely to lead to a sequence restoration event that corrects the complex mCherry^fs-2;Y77X^ mutation, and thus cells with such an indel event for sequencing were not selected.

When comparing the editing events generated by Cas9 nuclease/AS-gRNA-treated cells relative to Cas9(D10A) nickase/AS-gRNA-treated cells, we note that there are more WT alleles generated when the Cas9 nuclease is used. However, we also noted lower coverage across the ONT reads when using the Cas9 nuclease (Figure S4A,C). Application of the Cas9(D10A) nickase, however, leads to comparable levels of wild-type reads but no sequence coverage loss (Figure S4B,D). This suggests utilising a Cas9 nickase for allele conversion, where allele conversion potential may be retained without the caveat of compromised genomic integrity (Regan et al., 2025), may be better than use of a Cas9 nuclease.

Further evidence in support of allele conversion events can be seen in detailed analysis of the flow cytometry data in unsorted cells after transfection with AS-gRNAs and Cas9 nuclease or Cas9(D10A) nickase in Figure 3B. In an individual unedited CHACR cell, the green fluorescence is derived from the G allele only. In cells transfected with the mCherry^fs-2;Y77X^ AS-gRNA and Cas9 or Cas9 nickase, a single population of mCherry fluorescent cells with a linear relationship to EGFP fluorescence can be observed (Figure 3B). This linearity is because the EGFP and mCherry genes are driven by the same promoter, therefore expression is coupled. In contrast, in cells transfected with the EGFP^Δ4^ AS-gRNA, two distinct populations of mCherry fluorescent cells are evident, both with a linear relationship to EGFP fluorescence. In this scenario, the EGFP fluorescence can be derived from either the basally functional G allele or the repaired B allele, leading to an extra allele capable of EGFP expression per allele capable of mCherry expression. These two populations can be quantified as EGFP-high or EGFP-low expression (Figure 3C), depending on how many repaired alleles a cell might have. This demonstrates that while targeting either allele with AS-gRNAs can lead to comparable levels of allele conversion as measured by mCherry expression, which AS-gRNA is used will determine which allele is the donor and recipient, resulting in different phenotypic outcomes as measured by the EGFP fluorescent subpopulation.

### Base Editor Related DNA Nicking Can Trigger Allele Conversion

Having shown that a Cas9(D10A) nickase can mediate high levels of allele conversion, we wondered if an adenine base editor may have the potential to mediate allele conversion events, given that targeted DNA nicking is an instrumental step in the base editing process. To test this, we transfected plasmids encoding the adenine base editor ABE8e with the mCherry^fs-2;Y77X^ AS-gRNA into CHACR cells and screened for mCherry fluorescent cells (Figure 4A). Nicking the mCherry^fs-2;Y77X^ mutation on the G allele using either Cas9(D10A) or ABE8e resulted in mCherry fluorescent cells, albeit less so with ABE8e (Figure 4B).

**Figure 4.**
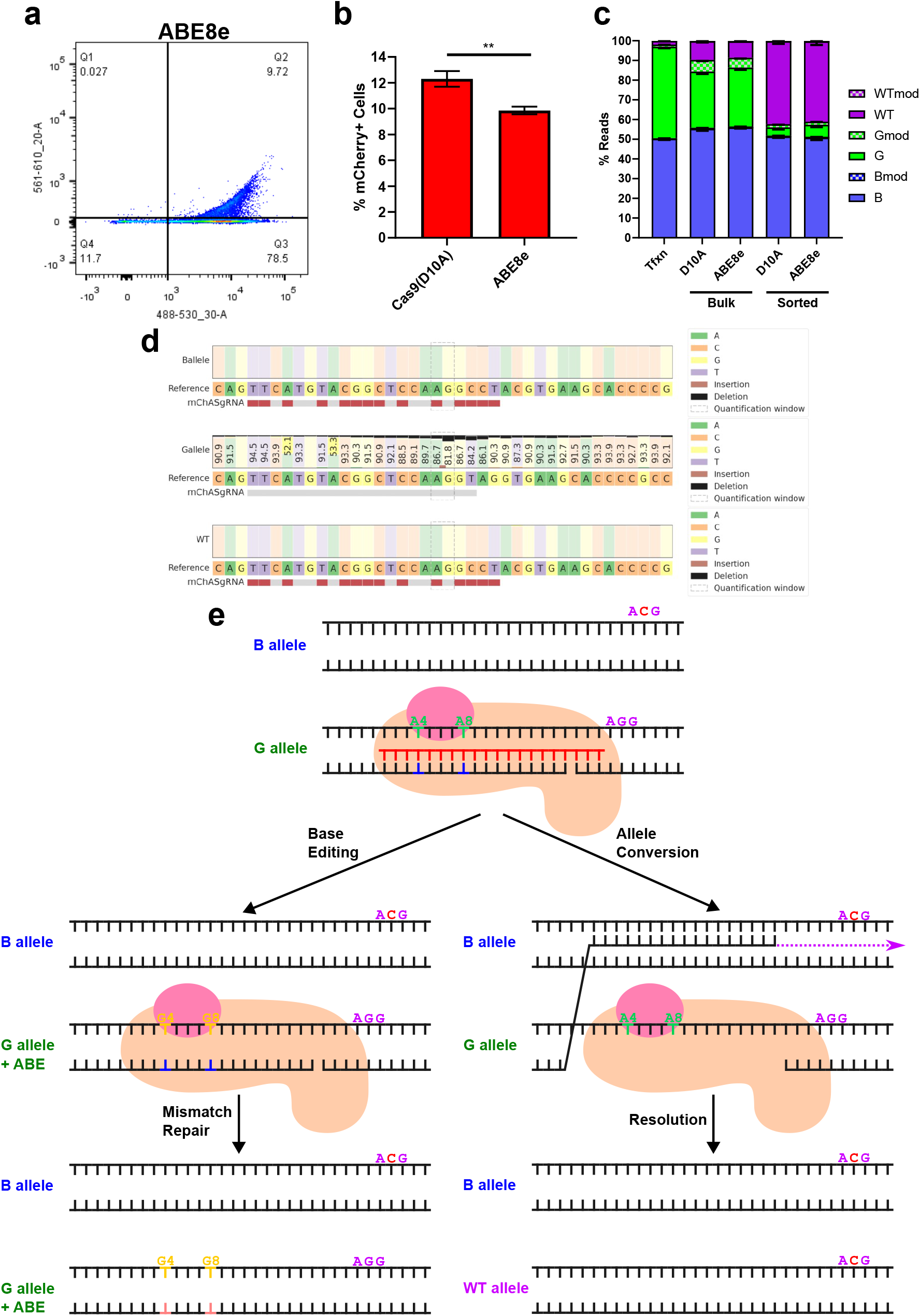
(a) Dot plot from CHACR cells transfected with ABE8e and mCherry AS-gRNA. (b) Quantification of mCherry fluorescent cells from CHACR cells transfected with either Cas9(D10A) or ABE8e with mCherry AS-gRNA. (c) ONT sequencing data for unsorted bulk populations or mCherry fluorescent sorted populations of CHACR cells transfected with either Cas9(D10A) or ABE8e with mCherry AS-gRNA across the EGFP and mCherry cassette. Sequencing data was analysed by CRISPResso2 for allele sequences matching either the B allele, G allele or a potential WT allele. (d) CRISPResso2 allele percentage quilts around the site of the mCherry AS-gRNA spacer sequence for each allele, from CHACR cells sorted for mCherry fluorescence after transfection with ABE8e and the mCherry AS-gRNA. Dotted box indicates the nicking window for ABE8e. (e) Hypothetical model for base editing/allele conversion decision point after the nick is created by ABE8e. Quantification of GFP^hi^ and GFP_lo_ cells as subsets of the total mCherry fluorescent population after allele conversion. Bars represent mean values from n=3 with error bars ± SD. * = *p*<0.05, ** = *p*<0.01, *** = *p*<0.001, **** = *p*<0.0001.

To characterise the molecular basis of the rescue of mCherry expression in cells transfected with ABE8e and the mCherry^fs-2;Y77X^ AS-gRNA, we performed long-read ONT sequencing of PCR products that span across the EGFP and mCherry cassette from genomic DNA harvested from either unsorted cells, or the mCherry fluorescent population of cells (Figure 4A, Q2). Indeed, the sorted mCherry fluorescent cells represent those with wild-type alleles, converted from the G allele by targeting the mCherry^fs-2;Y77X^ mutation, while the proportion of B allele reads is unchanged (Figure 4C).

To determine if any canonical base editing events mediated by ABE8e and the mCherry^fs-2;Y77X^ AS-gRNA had occurred in these cells, both the bulk and mCherry sorted populations were analysed. Only reads ranked as the G allele contained adenine base editing at position 4 and 8, which is within the expected editing window for ABE8e (Richter et al., 2020). Furthermore, the wild-type allele reads in allele converted cells most likely derive from the G allele, as the G allele is specifically targeted by the mCherry^fs-2;Y77X^ AS-gRNA and the frequency of G allele reads drops in mCherry fluorescent cells making way for WT allele reads. However, while ABE8e is triggering allele conversion of G allele reads into wild-type allele reads, adenine base editing was not evident in wild-type allele reads (Figure 4D). This data indicates that a decision point occurs after allele-specific DNA nicking by ABE8e in heterozygous cells, where either allele conversion or base editing can occur, but base edits and allele conversion information are not combined into a unique sequence (Figure 4E).

### Allele Conversion Efficiency Can Be Modified by Manipulating RAD51 and DNA-PKcs

We next investigated potential molecular mechanisms that may modify the allele conversion events observed in this model. We decided on three approaches: first to transfect with plasmids encoding gene editing modifiers; second, to transfect with plasmids encoding additional modifier gRNAs; and third, to test with small molecule inhibitors also known to affect gene editing efficiency.

We first started by transfecting CHACR cells with Cas9(D10A) and the mCherry^fs-2;Y77X^ AS-gRNA in combination with plasmids encoding common modifiers of gene editing and DNA repair. A dominant negative variant of the mismatch repair protein MLH1 is often applied in prime editing systems (PE4/5) to enhance editing efficiency (Chen et al., 2021). We hypothesised that mismatch repair mechanisms may be engaged to resolve the information copied from the heterologous allele. Here, we observed a small decrease in allele conversion efficiency as evident by reduced mCherry fluorescent cells when cotransfecting with dnMLH1 (Figure 5A,B). Next, we overexpressed human RAD51 and the dominant negative variant RAD51(K133R), which have been shown to decrease and increase HDR efficiency, respectively (Rees et al., 2019). Overexpression of either wild-type or K133R mutant RAD51 both reduced allele conversion efficiency, suggesting a complex function of this component in this system. Finally, we overexpressed a ubiquitin variant that selectively binds and inhibits 53BP1 (i53), which has also been shown to improve HDR (Canny et al., 2018). Again, we observed a small decrease in allele conversion efficiency, implicating a role for 53BP1 in the allele conversion pathway.

**Figure 5.**
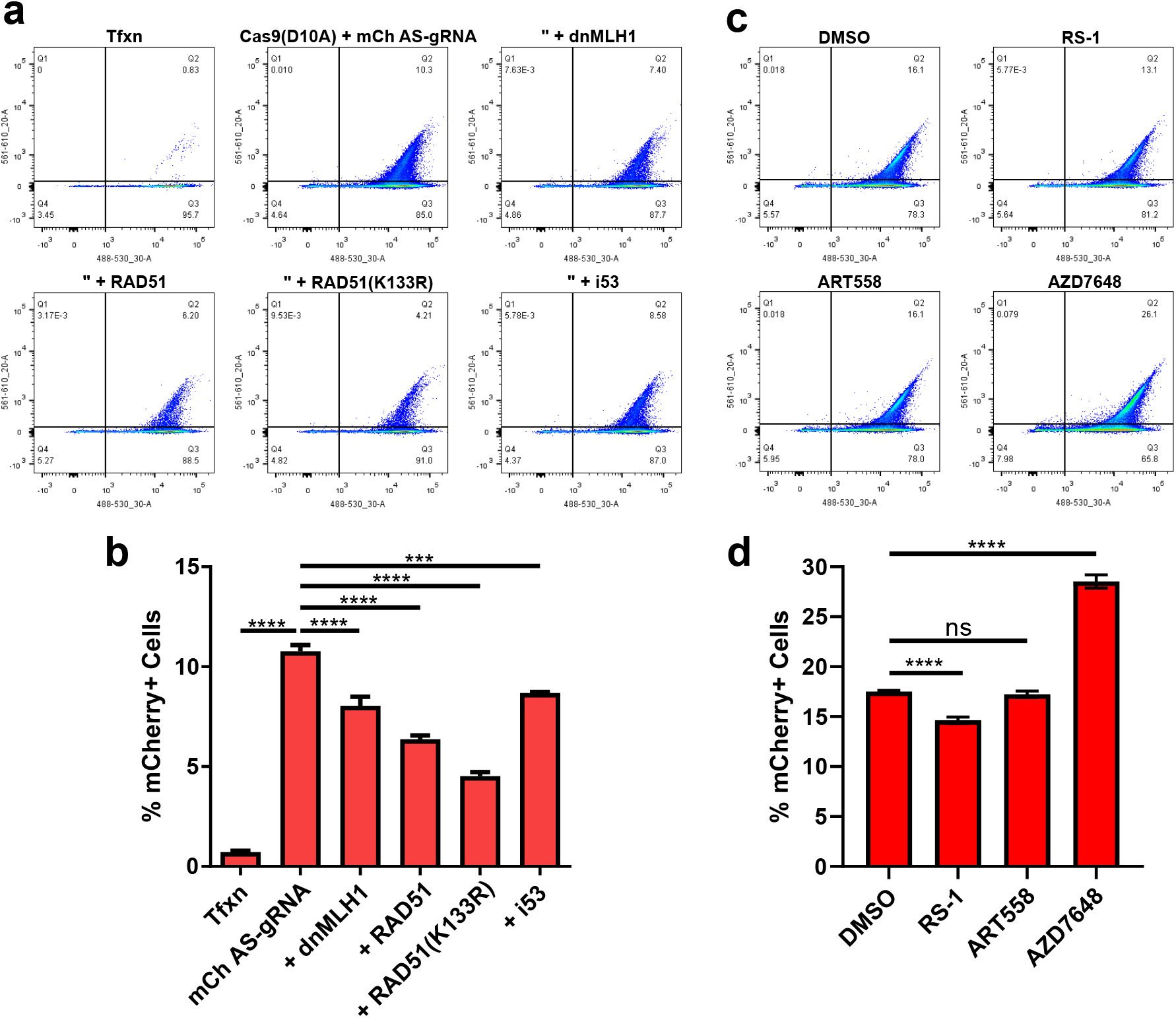
(a) Dot plots from transfection control CHACR cells, or transfected with Cas9(D10A) and mCherry AS-gRNA, as well as plasmids expressing editing modifiers. (b) Accompanying quantification of mCherry fluorescent cells after allele conversion with editing modifiers. (c) Dot plots from CHACR cells transfected with Cas9(D10A) and mCherry AS-gRNA treated with editing modifying compounds. (d) Accompanying quantification of mCherry fluorescent cells after allele conversion with modifying compound treatment. Bars represent mean values from n=3 with error bars ± SD. * = *p*<0.05, ** = *p*<0.01, *** = *p*<0.001, **** = *p*<0.0001.

In another model of allele conversion, interhomolog recombination was triggered by coadministration of an AS-gRNA targeting a gene-inactivating mutation as well as a secondary, allele-agnostic gRNA (AA-gRNA) (Tomita et al., 2023). This AA-gRNA was shown to have the potential to improve allele conversion when compared to AS-gRNA alone. Thus, we designed 6 AA-gRNAs targeting across the fluorescence genes (Figure S5A) and co-transfected these as plasmids with the mCherry^fs-2;Y77X^ AS-gRNA and Cas9(D10A). The addition of these secondary AA-gRNAs reduced the amount of mCherry fluorescent cells (Figure S5B), indicating a reduction of allele conversion efficiency in the CHACR cell model.

Small molecule compounds have also been developed with the aim of improving gene editing outcomes. The RAD51 stimulatory compound RS-1 has been shown to improve HDR editing efficiency (Song et al., 2016), however we observed a slight reduction in allele conversion efficiency (Figure 5C,D), echoing the outcomes of RAD51 overexpression. POLQ, which is inhibited by ART588, has been shown to be involved in interhomolog recombination that can lead to spontaneous gene correction at sites of DNA damage (Davis et al., 2020). Despite this, ART558 treatment led to no difference in mCherry fluorescent cells compared to DMSO vehicle control. Finally, DNA-PKcs have been similarly implicated in interhomolog recombination (Davis et al., 2020), and inhibition of DNA-PKcs using AZD7648 has been shown to improve HDR efficiency (Wimberger et al., 2023). AZD7648 treatment significantly increased mCherry fluorescent cells compared to DMSO vehicle control, indicating an inhibitory role for DNA-PKcs in allele conversion.

## Discussion

Emerging gene editing technologies have offered unprecedented potential for the correction of genetic variants underlying a plethora of diseases. With the increasing complexity of editing required, more intricate editing techniques are required leading to cumbersome cargoes and complicated designs. Here, we investigated an allele conversion mechanism that may allow for the correction of heterozygous variants as an alternative to current editing strategies. To investigate allele conversion, we produced the CHACR cell line. This model includes compound heterozygous variants across genes encoding fluorescent proteins stably integrated into the genome. Allele conversion was triggered using a Cas9 nuclease or nickase and a gRNA specifically targeting one of the heterozygous mutations inactivating mCherry fluorescence. Application of Cas9 with an AS-gRNA targeting either the B or G allele led to recovery of mCherry fluorescence, and this fluorescent population was subsequently sorted and sequenced. This revealed that mCherry fluorescent cells harboured fully wild-type sequences, indicating that an allele conversion event corrected the fluorescence-inactivating mutations.

As a byproduct, we also detected modified versions of the B or G allele reads. Modified reads likely arise from a Cas9-associated indel that best matches either the B or G allele amplicon. While the Cas9 nuclease and Cas9(D10A) nickase conditions seem to be allele and sequence specific, there seems to be no discernible difference in the amount of modified reads resulting from either the nuclease or nickase. In agreement with this, the most common modified read from these conditions is a 1nt insertion on the targeted allele, immediately upstream of the expected cut site (Figure S2A-D), has been shown to be a frequent outcome of Cas9-induced DSBs (Lemos et al., 2018). Importantly, because we observed allele conversion with either Cas9 nuclease or nickase, we tested these components at the *HEK3* control locus (Figure S6A) and confirmed that the Cas9 nuclease (Figure S6B) but not the Cas9(D10A) nickase (Figure S6C) was producing indels at a control locus. This suggests that either a DSB or nick can trigger allele conversion, or that an allele specific nick at a heterozygous site can be converted to a DSB as part of the allele conversion process.

Upon targeted allele conversion to correct either the EGFP^Δ4^ frameshift on the B allele or the complex mCherry^fs-2;Y77X^ mutation on the G allele, CHACR cells produced measurable mCherry fluorescence. The linear correlation between EGFP and mCherry fluorescence intensity after allele conversion demonstrated that these proteins were expressed from the same transcript. Long read ONT sequencing confirmed that the EGFP and mCherry wild-type sequences that once existed exclusively on separate chromosomes occurred on the same sequencing read. This suggests a type of crossover typically observed during meiotic recombination, where information is copied between the homologous chromosomes after DNA damage incurred by SPO11 (Zheng et al., 2025). Here, a Cas9 nuclease or nickase maybe mimicking the role of SPO11, followed by DNA repair machinery triggering allele conversion.

This allele conversion event orchestrated by allele-specific targeting of Cas9 has been observed by others. Interhomolog repair (IHR) can be triggered by Cas9 nuclease or nickase in a cell line derived from HT1080 fibrosarcoma cells at a very low frequency to recover CD44 expression (Davis et al., 2020). A similar outcome was observed using AS-gRNAs in *Drosophila* (Roy et al., 2022) and *Arabidopsis* (Zhang et al., 2025), suggesting an evolutionarily conserved mechanism engaged outside of meiosis. Tomita et al. showed that nickase-triggered allele conversion is extremely low in TK6261 lymphoblast cells, but is enhanced by the application of an additional, allele-agnostic gRNA (Tomita et al., 2023). Similarly, IHR was observed to be a rare event that followed DSBs, which can be amplified by depleting the BLM helicase (LaRocque et al., 2011, Regan et al., 2025). The higher frequency of allele conversion observed in our CHACR cells may be due to their HEK293T-based ancestry, which have known DNA repair deficiencies (Trojan et al., 2002) that may point towards an allele conversion mechanism.

The administration of an extra gRNA to improve IHR as per NICER (Tomita et al., 2023) did not increase the amount of mCherry fluorescent cells our CHACR system, which may also be explained by overburdened DNA repair mechanisms. In analogy, prime editing efficiency has been shown to increase when adding synonymous variants across the reverse transcription template to bias mismatch repair towards the edit (Chen et al., 2021). The secondary AA-gRNA may bias IHR towards allele conversion in the NICER system by increasing DNA damage burden, while the DNA repair deficient HEK293T-derived CHACR system may only require the sole AS-gRNA for allele conversion. Furthermore, an additional gRNA may quench the Cas9 leading to reduced efficiency, which may also explain the reduced efficiency in the CHACR system when applying extra gRNAs. If this is the case, modulating DNA repair mechanisms may be a more effective avenue for improving allele conversion efficiency than using an AA-gRNA.

Finally, we aimed to apply the CHACR model to investigate the mechanism behind allele conversion by manipulating DNA repair mechanisms. Mismatch repair inhibition has been shown to improve prime editing efficiency (Chen et al., 2021), and we hypothesised a similar improvement in allele conversion considering that there is likely a similar intermediate DNA heteroduplex between the recipient and donor alleles. Conversely, dnMLH1 transfection reduced the amount of mCherry fluorescent CHACR cells, suggesting mismatch repair inhibition reduced allele conversion efficiency. This may be due to the existing mismatch repair deficiency in HEK293T cells (Panigrahi et al., 2012), or that a repair mechanism that requires functional MLH1 is engaged for allele conversion.

We also showed a requirement for RAD51 in allele conversion, where the biggest reduction in mCherry fluorescent cells was observed when applying the dominant-negative RAD51(K133R) mutant (Rees et al., 2019). However, overexpression of wild-type RAD51 and stimulation of RAD51 with RS-1 also led to a reduction in allele conversion efficiency. Further work must be carried out to decipher the role of RAD51 in allele conversion and how to manipulate strand invasion to improve efficiency. Importantly, we showed that DNA-PKcs inhibition using AZD7648 noticeably increased allele conversion efficiency. This finding echoes the effect of NU7441, another DNA-PKcs inhibitor, which was shown to improve IHR efficiency at the site of DSBs (Davis et al., 2020). Additionally, we showed that DNA-PKcs inhibition was able to increase allele conversion at the site of an allele specific nick rather than a DSB. Considering that DNA-PKcs canonically recognise DSBs to trigger non-homologous end joining (NHEJ), this further suggests that allele specific nicks may be converted to DSBs before allele conversion. Alternatively, DNA-PKcs may recognise these allele specific nicks in a non-canonical mechanism antagonistic to allele conversion.

The application of AS-gRNAs to trigger allele conversion has been observed in disease relevant models, introducing a precedent for a therapeutic niche. Targeting the sickle cell disease related gene *HBB* with a Cas9 nuclease was shown to introduce sequences from the *HBD* gene, indicating gene conversion between the two loci with curative potential (Javidi-Parsijani et al., 2020). Recently, allele conversion was achieved in leukaemia cell models and primary haematopoietic cells using a Cas9 nuclease and AS-gRNAs targeting cancer associated heterozygous mutations (Silver et al., 2025). Here, in agreement with previous reports (Roy et al., 2022, Tomita et al., 2023), we show that the corrective potential of allele conversion can be instigated with a Cas9 nickase, thereby limiting the risk of indels potentially worsening the target sequence context when using a nuclease (Regan et al., 2025). To advance nickase-based allele conversion as a strategy for correcting pathogenic heterozygous mutations in recessive or dominant genetic disease, we endeavour to apply the CHACR system to further identify the mechanism behind allele conversion and improve efficiency.

## Materials and Methods

### Cell Culture

HEK293T cells were maintained in DMEM high glucose, GlutaMAX (Gibco) supplemented with 10% fetal bovine serum. Cells were transfected with plasmids using Lipofectamine 3000 (Invitrogen) by seeding 300,000 cells per well in a 6 well plate the day prior to transfection. On the day of transfection, Lipofectamine 3000 was prepared in OptiMEM (Gibco) as per the manufacturer protocol with 2µg total plasmid DNA. For allele conversion experiments, plasmids were transfected at a 3:1 ratio of Cas9:gRNA. hCas9 (Addgene #41815) and hCas9(D10A) (Addgene #41816) were kind gifts from George Church. ABE8e (Addgene #138489) was a kind gift from David Liu. Drug compounds were sourced from MedChemExpress and resuspended in DMSO, and dilutions were applied to cells shortly before transfection with Cas9(D10A) and gRNA plasmids.

### gRNA Cloning, Transformation, and Propagation

gRNA spacer sequences were cloned into the gRNA Cloning Vector Bbs I ver. 2 (Addgene #85586, a kind gift from Hodaka Fujii). 20nt spacer sequences (Additional Information) were ordered as short top and bottom complementary oligonucleotides from Integrated DNA Technologies (IDT EU) with complementary overhangs to BbsI-digested gRNA Cloning Vector plasmids. gRNA cloning into this vector was conducted by Golden Gate assembly. Resulting gRNA plasmids were transformed into NEB® 5-alpha Competent *E. coli* (High Efficiency) by heat shock: incubation with gRNA plasmid for 30min on ice, heat shock at 42°C for 30s, followed by incubation on ice for 5min. Cultures were recovered with SOC medium at 37°C for 1h followed by spreading onto ampicillin-supplemented LB agar plates and incubated at 37°C. The following morning, colonies were picked and propagated in 5mL of LB medium supplemented with ampicillin, followed by plasmid extraction using NucleoSpin™ Plasmid DNA (Fisher). Resulting plasmid preparations were sequenced using Plasmidsaurus plasmid sequencing service, and positive colonies were propagated in larger 50mL cultures for higher yield plasmid extraction using Plasmid *Plus* Midi Kit (Qiagen).

### CHACR Model Design and Development

Knock-in plasmids expressing B allele and G allele constructs were designed and produced using VectorBuilder. These plasmids were transfected into HEK293T cells using Lipofectamine 3000 with pSpCas9(BB)-2A-Puro (PX459) V2.0 (Addgene #62988, a kind gift from Feng Zhang) and a plasmid expressing a gRNA targeting the *AAVS1* locus. 72h after transfection, cells were treated with 1µg/mL puromycin for a week, until separate control cell cultures were dead. Puromycin resistant cells were recovered and subject to FACS for EGFP+, mCherry- and mTagBFP2+ cells and sorted for single cells per well into 96 well plates. Resulting colonies were expanded, and DNA samples were extracted using QuickExtract DNA Extraction Solution (LGC) as per manufacturer protocol. Screening for positive clones was conducted by PCR amplification of the endogenous *AAVS1* locus, the 5’ junction between *AAVS1* locus and knock-in cassette, and the EGFP-mCherry cassette for genotyping. After first attempt, a resulting clone only contained knock-in of the G allele, so this clone was used for a second knock-in process using a gRNA targeting the mCherry cassette and the B allele knock-in plasmid. This led to the final CHACR clone developing a 2nt deletion in the mCherry cassette of the G allele which is not found on the knock-in plasmid. The final CHACR clone was propagated and stocked, and a stable EGFP fluorescence profile was confirmed >6 months after the original knock-in transfection.

### Flow Cytometry and Cell Sorting

In preparation for flow cytometry, cells were dissociated from culture vessels using TrypLE Express Enzyme (Gibco) and pelleted at 200xg for 5min. Medium supernatant was aspirated, and pellets were resuspended in phosphate buffered saline without calcium and magnesium and dispensed into flow cytometry tubes through cell strainers (Falcon). Resulting cell suspensions were analysed and/or sorted using BD FACSAria Fusion with filters for EGFP (488nm excitation, 530nm filter), mCherry (561nm excitation, 610nm filter) and mTagBFP2 (405nm excitation, 450nm filter). After sorting, cells were dispensed into either 96 well plates for single cell sorting, or into 15mL tubes with DMEM supplemented with 10% FBS for mCherry fluorescent cell sequencing after allele conversion experiments. For post-allele conversion sequencing, treatment conditions that produced at least 10% mCherry fluorescent cells were eligible for sorting, as lower yields would not produce enough cells for immediate DNA extraction.

### Polymerase Chain Reaction and Purification

Genomic DNA extracts were used as template for PCRs to amplify target sequences using Q5 Hot Start Polymerase (NEB). PCR reactions were prepared as per manufacturer protocol using forward and reverse primers produced as oligonucleotides from IDT (Additional Information). A sample of PCR products were resolved on a 1% agarose gel at 100V to confirm successful amplification, and the remaining PCR product was purified using Sera-Mag SpeedBeads (Merck) at a 1:1 ratio by volume. During purification, PCR product and magnetic bead mixtures were washed twice with freshly prepared 80% ethanol before eluting in nuclease free water. Purified PCR products were quantified using Qubit dsDNA BR Assay and diluted to concentrations required for next generation sequencing (NGS). Illumina NGS was conducted using Genewiz Amplicon-EZ service. Long read ONT sequencing was conducted using Plasmidsaurus Premium PCR Sequencing service.

### Data Analysis

Resulting NGS.fastq files were analysed CRISPResso2. Long-read sequences produced from genotyping the CHACR clone were aligned to the human genome with a synthetic EGFP-mCherry cassette appended to allow for visualisation of the EGFP and mCherry mutations in Integrated Genome Viewer (IGV). Amplicon sequences for the G allele, B allele and expected WT allele were assessed for allele frequency quantification. Flow cytometry data was analysing using FlowJo V10. Data was graphed and analysed using GraphPad Prism 8.3.

