## Supplemental Figure 1 for "Interrogating the Mechanisms of Cas9-mediated Allele Conversion"

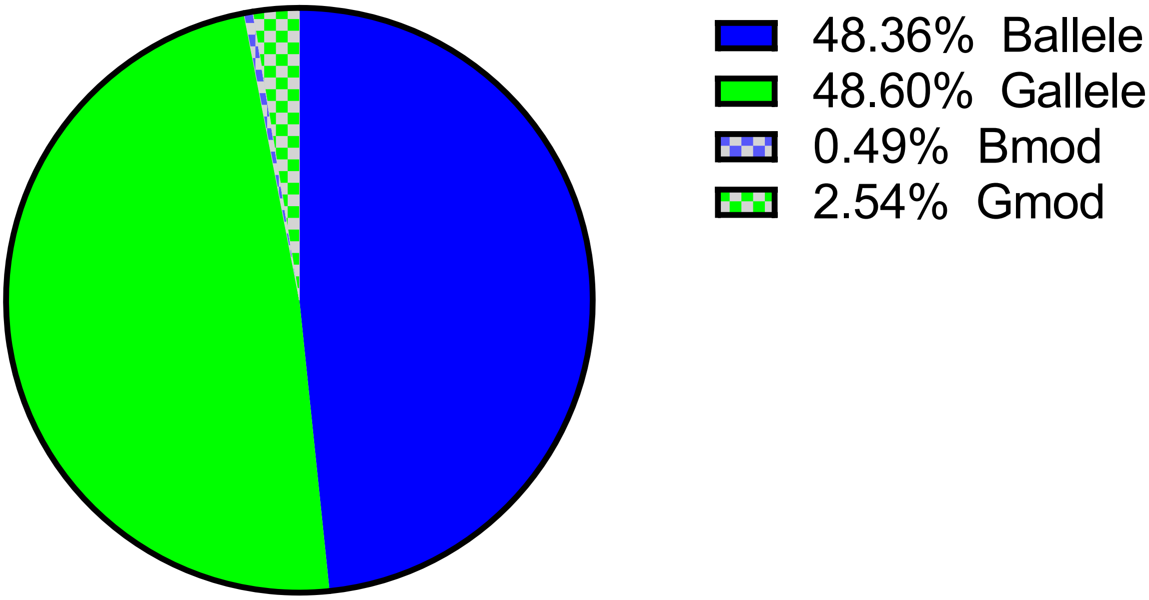


Figure S1: Proportions of alleles in the final CHACR cell line. Long read ONT sequencing was analysed by CRISPResso2 command with inputs for the expected sequences of the B allele and G allele spanning the EGFP to mCherry cassettes. Modified reads include sequences that align best with a specific allele including indels and/or substitutions.

Table S1:Top two highest represented reads for Illumina NGS of the mTagBFP2 cassette of a candidate reporter clone. The 2nt insertion in the G allele mTagBFP2 is highlighted in yellow. Illumina .fastq analysis performed using Azenta UniqueSeq. Next highest represented read accounted for 0.3% of total reads, which was considered sequencing error. Sequencing of this clone suggested potential tetraploidy of the parental line, therefore CHACR cells may have four copies of the *AAVS1* locus to integrate into.

| Sequence | Count | % |
|---|---|---|
| AGGGCGAAGGCAAGCCCTACGAGGGCACCCAGACCATGAGAATCAAGGTG<br>GTCGAGGGCGGCCCTCTCCCCTTCGCCTTCGACATCCTGGCTACTAGCTT<br>CCTCTACGGCAGCAAGACCTTCATCAACCACACCCAGGGCATCCCCGACT<br>TCTTCAAGCAGTCCTTCCCTGAGGGCTTCACATGGGAGAGAGTCACCACA<br>TACGAAGACGGGGGCGTGCTGACCGCTACCCAGGACACCAGCCTCCAGGA<br>CGGCTGCCTCATCTACAACGTCAAGATCAGAGGGGTGAACTTCACATCC | 34831 | 74.902 |
| AGGGCGAAGGCAAGCCCTACGAGGGCACCCAGACCATGAGAATCAAGGTG<br>GTCGAGGGCGGCCCTCTCCCCTTCGCCTTCGACATCCTGGCTACTAGCTT<br>CCTCTACGGCAGCAAGACCTGGTCATCAACCACACCCAGGGCATCCCCGA<br>CTTCTTCAAGCAGTCCTTCCCTGAGGGCTTCACATGGGAGAGAGTCACCA<br>CATACGAAGACGGGGGCGTGCTGACCGCTACCCAGGACACCAGCCTCCAG<br>GACGGCTGCCTCATCTACAACGTCAAGATCAGAGGGGTGAACTTCACATC<br>C | 11671 | 25.097 |

5F2E
