## Supplemental Figure 2 for "Interrogating the Mechanisms of Cas9-mediated Allele Conversion"

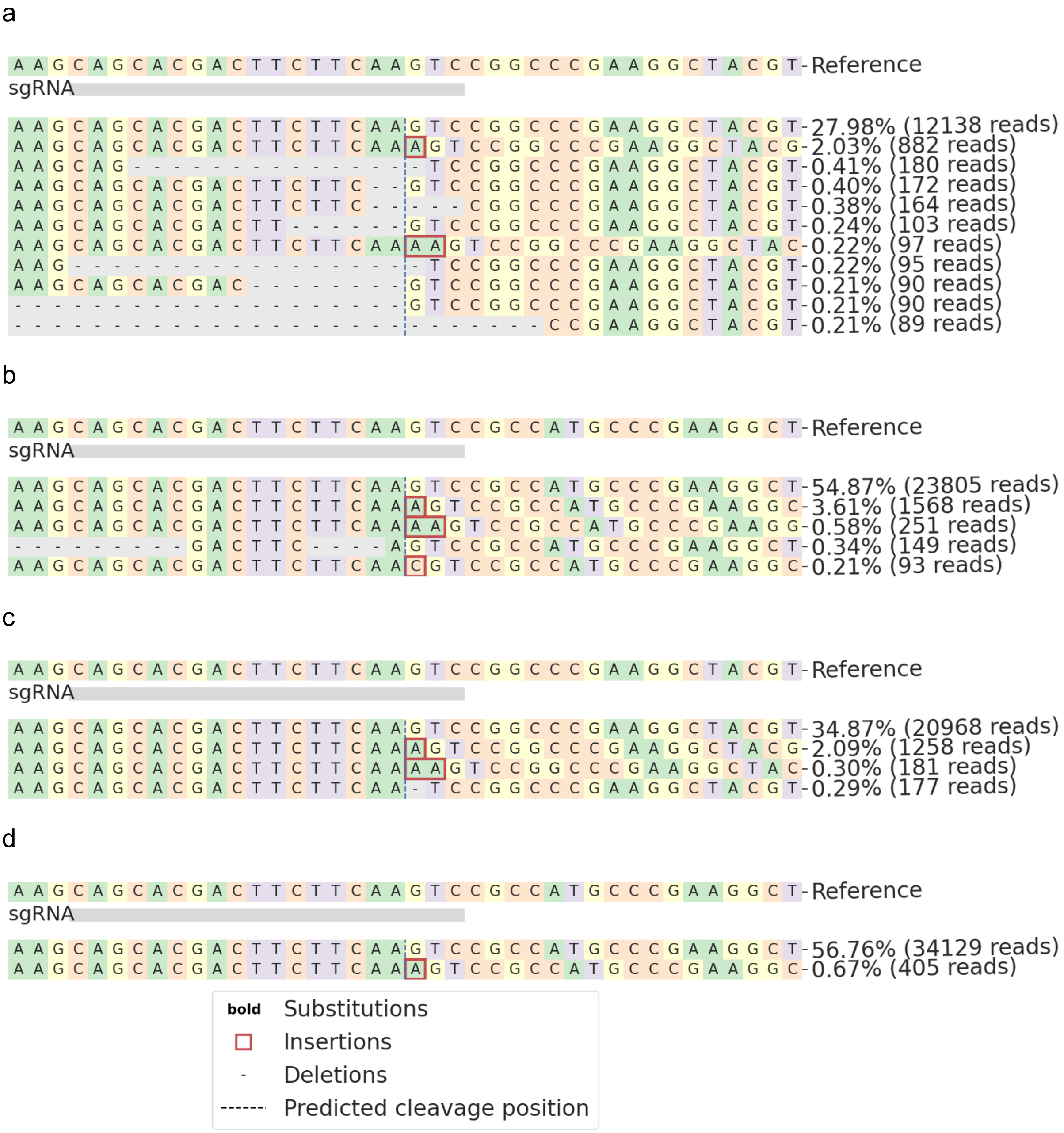


Figure S2: CRISPResso2 allele frequency plots around the EGFP$^{\Delta 4}$ AS-gRNA site in reporter cell on a PCR amplicon for the EGFP cassette. Allele frequency plots for cells transfected with Cas9 and the EGFP$^{\Delta 4}$ AS-gRNA around (A) the B allele or (B) the G allele. Allele frequency plots for cells transfected with Cas9(D10A) and the EGFP$^{\Delta 4}$ AS-gRNA around (C) the B allele or (D) the G allele. The most represented modified read is an adenine insertion at the cut site for both alleles, regardless of transfection with Cas9 or Cas9(D10A).
