## Supplemental Figure 3 for "Interrogating the Mechanisms of Cas9-mediated Allele Conversion"

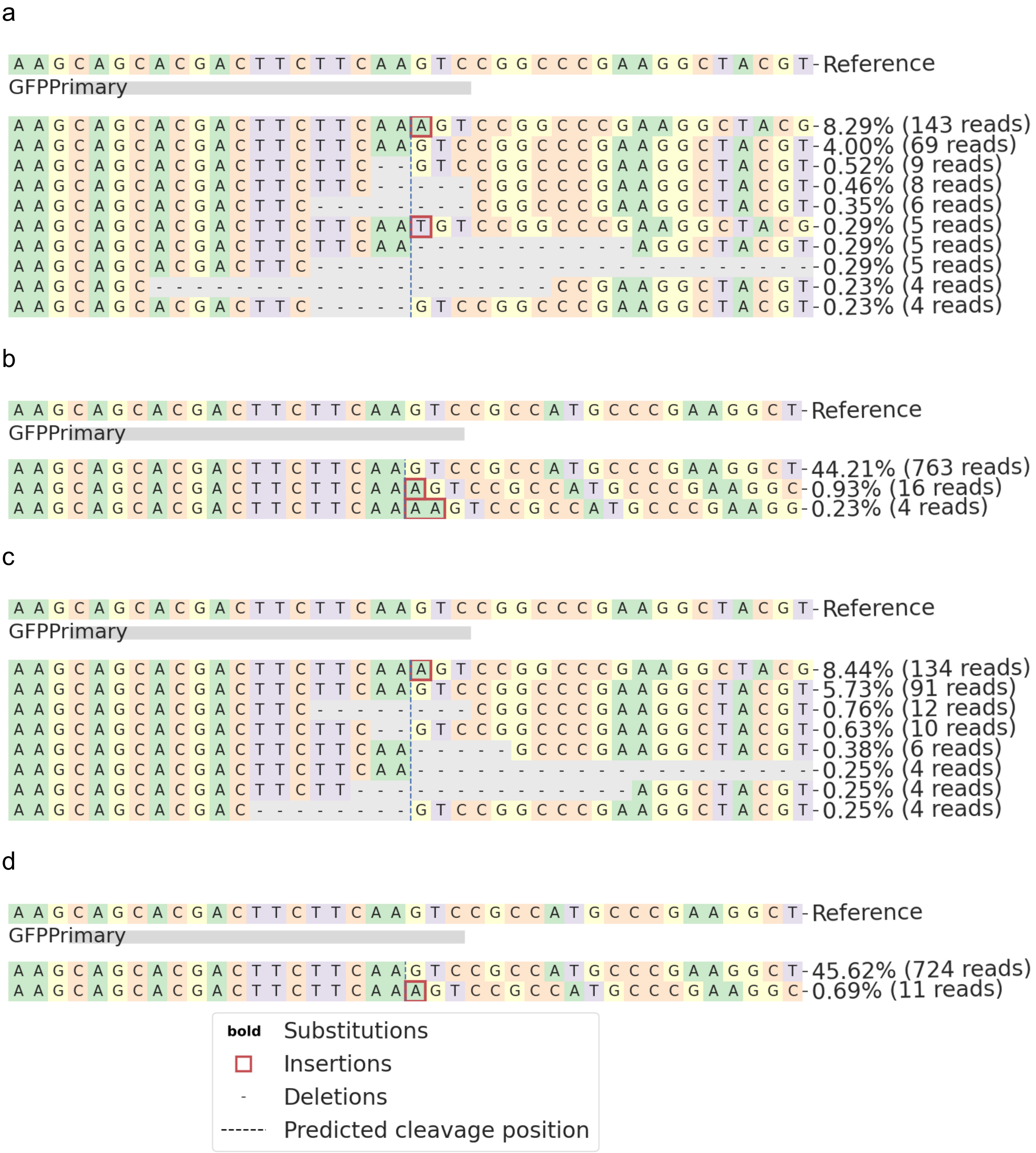

Figure S3: CRISPResso2 allele frequency plots around the EGFP AS-gRNA site in reporter cells sorted for mCherry fluorescence, on a PCR amplicon for EGFP to mCherry cassette. Allele frequency plots for cells transfected with Cas9 and the EGFP AS-gRNA around (A) the B allele or (B) the G allele. Allele frequency plots for cells transfected with Cas9(D10A) and the EGFP AS-gRNA around (C) the B allele or (D) the G allele. The most represented modified read is an adenine insertion at the cut site for both alleles, regardless of transfection with Cas9 or Cas9(D10A). Lower represented reads, such as deletions, are also predicted to restore the reading frame that will induce mCherry expression and fluorescence.
