## Supplemental Figure 4 for "Interrogating the Mechanisms of Cas9-mediated Allele Conversion"

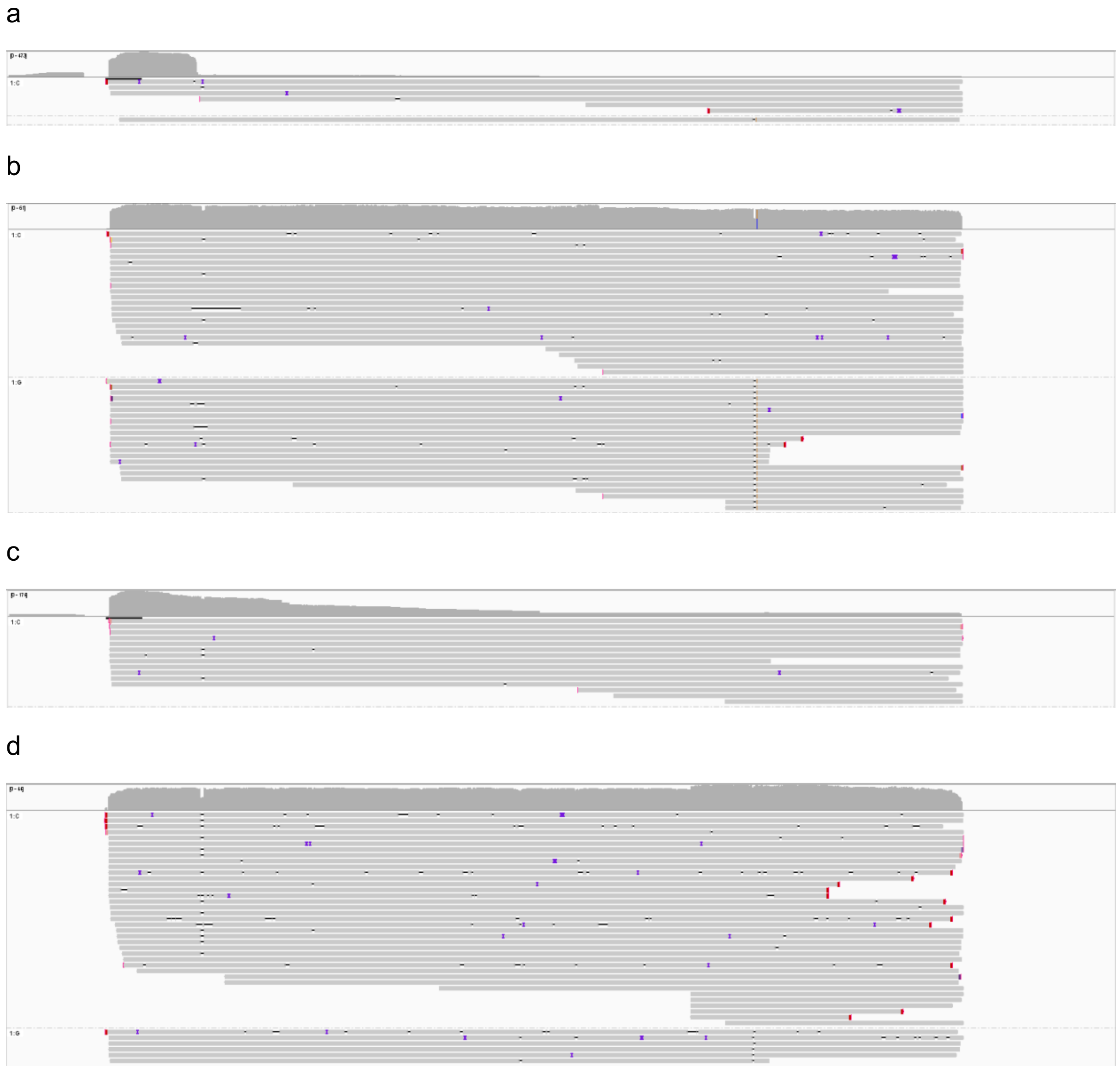

Figure S4: CHACR cells treated with (a) Cas9 or (b) Cas9(D10A) with EGFP AS-gRNA, and (c) Cas9 or (d) Cas9(D10A) with mCherry AS-gRNA were sorted for mCherry fluorescent cells can sequenced by long read ONT across the EGFP and mCherry cassettes. Resulting sequencing files were aligned against a synthetic, wild-type EGFP-mCherry sequence and visualised in IGV. Reads were grouped by the base at the PTC in the mCherry cassette to identify reads in that sample with a WT EGFP that phase with a WT mCherry. Reads in group 1: C have a wild-type mCherry sequence while reads in group 1: G have the mCherry PTC mutation.
