## Supplemental Figure 5 for "Interrogating the Mechanisms of Cas9-mediated Allele Conversion"

a

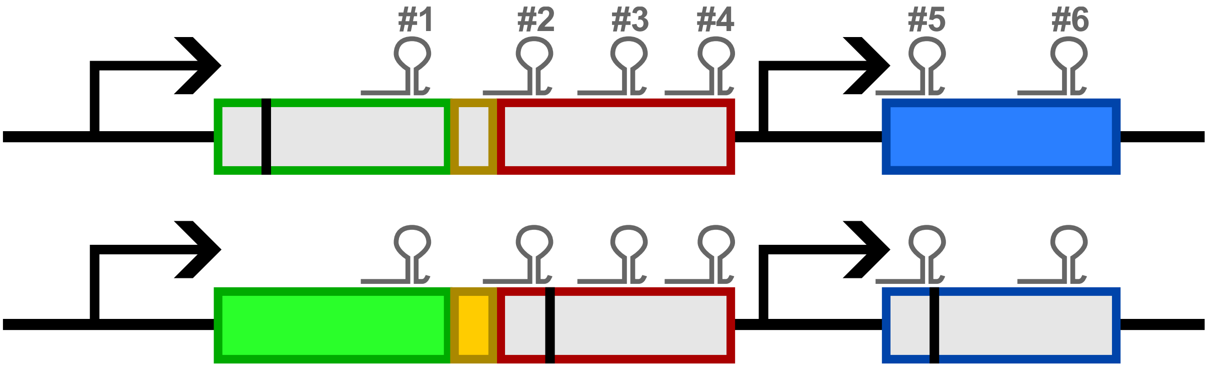


b

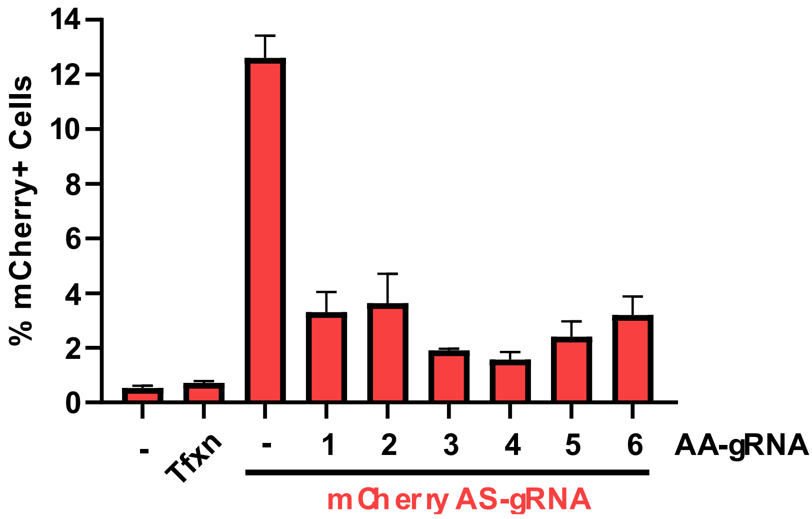


Figure S5: (A) Schematic of the locations of AA-gRNAs across the reporter cassette. (B) mCherry fluorescent CHACR cells percentage of the whole population 72h after transfection with plasmids encoding Cas9(D10A), the mCherry AS-gRNA with or without AA-gRNA #1-6. Data compared to untransfected CHACR cells or CHACR cells treated with the transfection reagent alone (Tfxn). Error bars represent mean ± SD from n=3.
