## Supplemental Figure 6 for "Interrogating the Mechanisms of Cas9-mediated Allele Conversion"

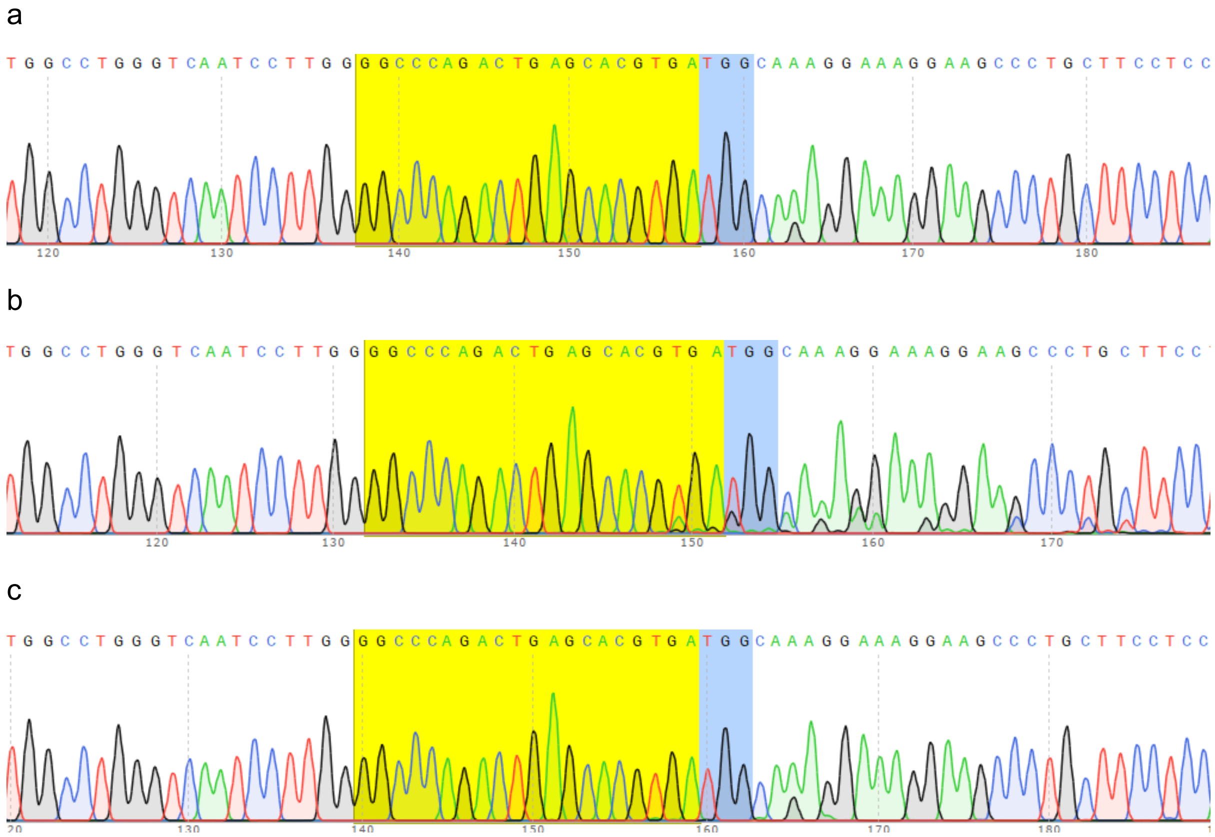


Figure 5: Sanger sequencing chromatograms of genomic DNA harvested from reporter cells at the HEK3 locus. (A) transfection control, (B) Cas9 nuclease and HEK3 targeting gRNA, (C) Cas9(D10A) and HEK3 targeting gRNA. The HEK3 targeting gRNA protospacer is highlighted in yellow and the Cas9 PAM is highlighted in blue. 3nt downstream of the PAM site is a known heterozygous SNP that exists in HEK293T cells.
